# *In vivo* detection of DNA secondary structures using Permanganate/S1 Footprinting with Direct Adapter Ligation and Sequencing (PDAL-Seq)

**DOI:** 10.1101/2023.11.28.569002

**Authors:** Angelika Lahnsteiner, Sarah J. C. Craig, Kaivan Kamali, Bernadette Weissensteiner, Barbara McGrath, Angela Risch, Kateryna D. Makova

## Abstract

DNA secondary structures are essential elements of the genomic landscape, playing a critical role in regulating various cellular processes. These structures refer to G-quadruplexes, cruciforms, Z-DNA or H-DNA structures, amongst others (collectively called ‘non-B DN’), which DNA molecules can adopt beyond the B conformation. DNA secondary structures have significant biological roles, and their landscape is dynamic and can rearrange due to various factors, including changes in cellular conditions, temperature, and DNA-binding proteins. Understanding this dynamic nature is crucial for unraveling their functions in cellular processes. Detecting DNA secondary structures remains a challenge. Conventional methods, such as gel electrophoresis and chemical probing, have limitations in terms of sensitivity and specificity. Emerging techniques, including next-generation sequencing and single-molecule approaches, offer promise but face challenges since these techniques are mostly limited to only one type of secondary structure. Here we describe an updated version of a technique permanganate/S1 nuclease footprinting, which uses potassium permanganate to trap single-stranded DNA regions as found in non-B structures, in combination with S1 nuclease digest and adapter ligation to detect genome-wide non-B formation. To overcome technical hurdles, we combined this method with direct adapter ligation and sequencing (PDAL-Seq). Furthermore, we established a user-friendly pipeline available on Galaxy to standardize PDAL-Seq data analysis. This optimized method allows the analysis of many types of DNA secondary structures that form in a living cell and will advance our knowledge of their roles in health and disease.

## 1. Introduction

The discovery of the double-helical structure of DNA by Watson and Crick (Watson & Crick, 1953) was a defining moment in science that revolutionized our understanding of genetics. However, beyond the well-known B conformation, DNA can adopt a diverse array of secondary or noncanonical (non-B) structures that play pivotal roles in various biological processes (reviewed in (Wang & Vasquez, 2023)). Thirteen percent of the human genome has the potential to form secondary structures which include G-quadruplexes (G4s), cruciforms, Z-DNA structures, H-DNA (triplex) structures, single-strand loop structures (polyA/Ts), as well as RNA:DNA hybrid structures (R-loops; Figure 1) (Guiblet et al., 2021). For instance, G4s, the most studied non-B structures, form from guanine-rich sequences that fold into stacked tetrads held together by hydrogen bonds. *In vitro* G4-sequencing identified over 700,000 potential G4-forming sites in the human genome (Chambers et al., 2015).

**Figure 1.**
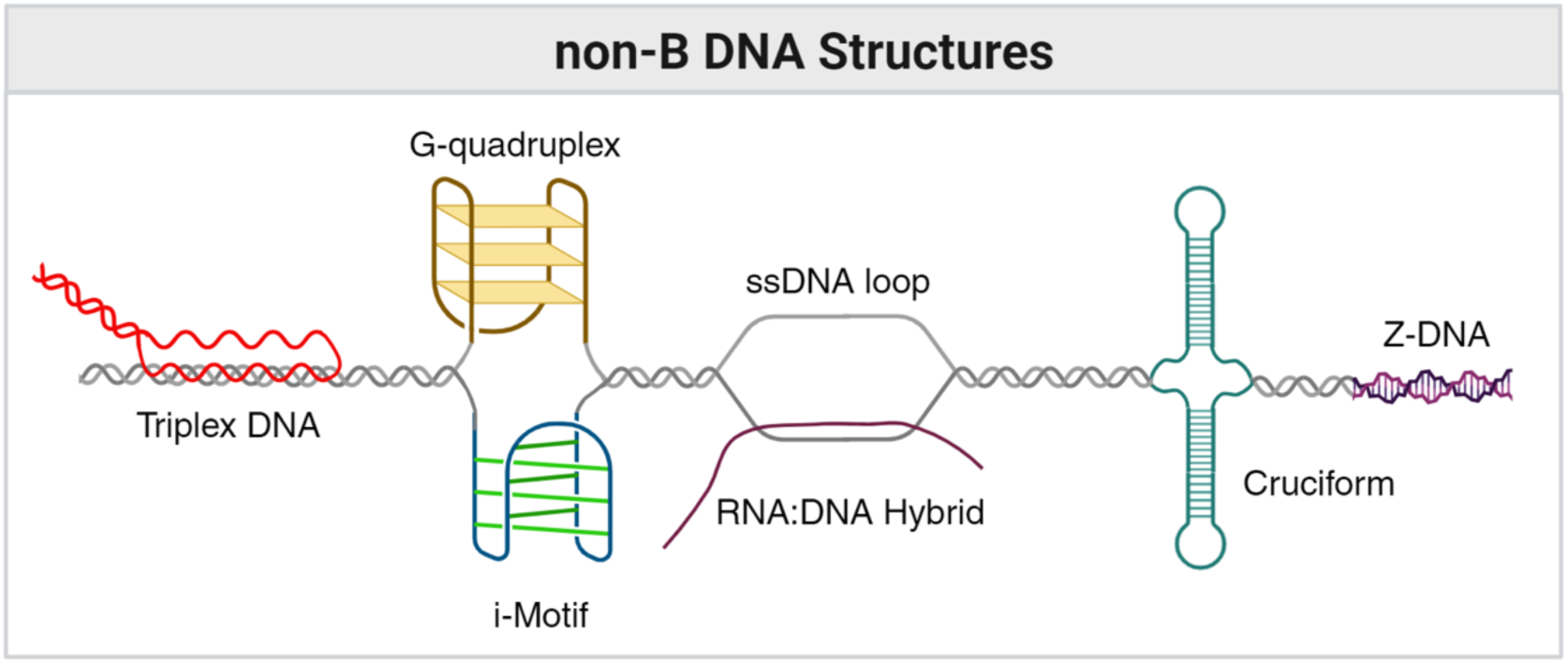
Non-B DNA structures. Various types of DNA repeats or motifs can form DNA secondary structures in living cells. Created with Biorender.com.

Initially, non-B DNA structures were described under mainly non-biological *in vitro* conditions. However, the latest highly specific and sensitive sequencing and microscopy techniques identified such structures *in vivo,* and revealed their diverse roles in many important cellular processes. For instance, they have been found to participate in recombination (Heissl et al., 2019; Kshirsagar, Khan, Joshi, Hosur, & Muniyappa, 2017; Saranathan, Biswas, Patra, & Vivekanandan, 2019), DNA repair (Brazda, Haronikova, Liao, Fridrichova, & Jagelska, 2016), transcription (Duquette, Handa, Vincent, Taylor, & Maizels, 2004; Georgakopoulos-Soares, Victorino, et al., 2022), epigenetics (Hansel-Hertsch et al., 2016; Sanz et al., 2016), determining cell identity and differentiation (Zyner et al., 2022), telomere maintenance (Chawla & Azzalin, 2008; Moye et al., 2015). Non-B structures have also been associated with mutations (Guiblet et al., 2021), DNA damage (De Magis et al., 2019), and cancer etiology (Hansel-Hertsch et al., 2020; Matos-Rodrigues et al., 2022) (also reviewed in (Bochman, Paeschke, & Zakian, 2012; Georgakopoulos-Soares, Chan, Ahituv, & Hemberg, 2022; Makova & Weissensteiner, 2023; Varshney, Spiegel, Zyner, Tannahill, & Balasubramanian, 2020)). This long list of biological processes reflects the importance of these structures in living cells (Wang & Vasquez, 2023), as well as in evolution (Makova & Weissensteiner, 2023), and highlights the need of reliable methods for their identification to facilitate our understanding of the underlying mechanisms.

DNA secondary structures can be detected with a wide range of methods, such as gel electrophoresis-based methods (Frank, Muller, & Wolff, 1981; Moon & Jarstfer, 2010), circular dichroism (CD) spectroscopy (Bishop & Chaires, 2003), atomic force microscopy (AFM) (Bose, Lech, Heddi, & Phan, 2018; Pyne & Hoogenboom, 2016), chromatin immunoprecipitation (ChIP) (Hansel-Hertsch et al., 2016; Hansel-Hertsch et al., 2020; Hansel-Hertsch, Spiegel, Marsico, Tannahill, & Balasubramanian, 2018; Sanz & Chedin, 2019; Sanz et al., 2016), permanganate/S1-footprinting (Kouzine et al., 2017; Kouzine et al., 2019), S1-Seq (Mimitou & Keeney, 2018), G4-Seq (Chambers et al., 2015), and polymerase stop assays (Wu & Han, 2019). Each of these techniques has its own strengths and limitations, and their choice often depends on the specific research question and the nature of the DNA secondary structures being studied. That each method tends to be specific for a single class of non-B structures, presents a significant impediment to global searches for all forms. This is especially true for the frequently used antibody-based ChIP-Seq techniques. Another primary concern with antibody-based methods is the specificity of the selected antibodies. Cross-reactions or non-specific binding events can lead to false positives. Furthermore, the performance of antibodies can vary from batch to batch. This variability can lead to inconsistent results and difficulties in reproducibility, making it challenging to compare data obtained from different laboratories and even experiments. Methods that are not based on antibodies have a greater potential to detect a broader variety of non-B DNA topologies simultaneously and thus are of great interest.

Potassium permanganate (KMnO_4_) footprinting has been widely used to map ssDNA formed by RNA polymerase binding to promoter regions (Bui, Rees, & Cotton, 2003; Craig et al., 1998; Davis, Bingman, Landick, Record, & Saecker, 2007; Gogos, Karayiorgou, Aburatani, & Kafatos, 1990), but more recently also to detect non-B DNA structures (Kouzine et al., 2013; Kouzine et al., 2017; Kouzine et al., 2019). There are several advantages of permanganate/S1 nuclease footprinting as opposed to several other methods for non-B detection based on S1 digests (Matos-Rodrigues et al., 2022; Mimitou & Keeney, 2018). KMnO_4_ preferentially and irreversibly oxidizes unpaired thymines under mild conditions compared to cytosine, guanine, and adenine which require harsher oxidation conditions (Akman, Doroshow, & Dizdaroglu, 1990). This fixation step traps the single-stranded state of the DNA and prevents reannealing or unfolding of non-B DNA during DNA extraction, especially, for instance, in regions of G4 formation. This provides a snapshot of the ssDNA-containing secondary structures in the cell, which would otherwise be lost during subsequent DNA preparation. Another important step of this method is that single stranded (SSB) or double stranded (DSB) DNA breaks, which occur naturally or result from DNA extractions, are blocked with terminal transferase. Blocking of these free ends avoids the detection of false-positive signals. A major drawback of the original versions of this method was that it required large amounts of the starting DNA material and therefore large amounts of enzymes and chemicals, making it rather expensive. More importantly, the very strong binding of the biotinylated fragments to the streptavidin coated beads led to poor recovery which greatly hindered the production of high-quality Illumina DNA sequencing libraries. To overcome this hurdle, we took advantage of permanganate/S1 nuclease footprinting published by Kouzine and colleagues (Kouzine et al., 2013; Kouzine et al., 2017; Kouzine et al., 2019), but updated the method to include direct Illumina adapter ligation as used in other methods such as S1-Seq (Mimitou & Keeney, 2018). We called the resulting technique *<u>P</u>ermanganate/S1 Nuclease Footprinting with <u>D</u>irect <u>A</u>dapter <u>L</u>igation and <u>Seq</u>uencing* (PDAL-Seq, Figure 2).

**Figure 2.**
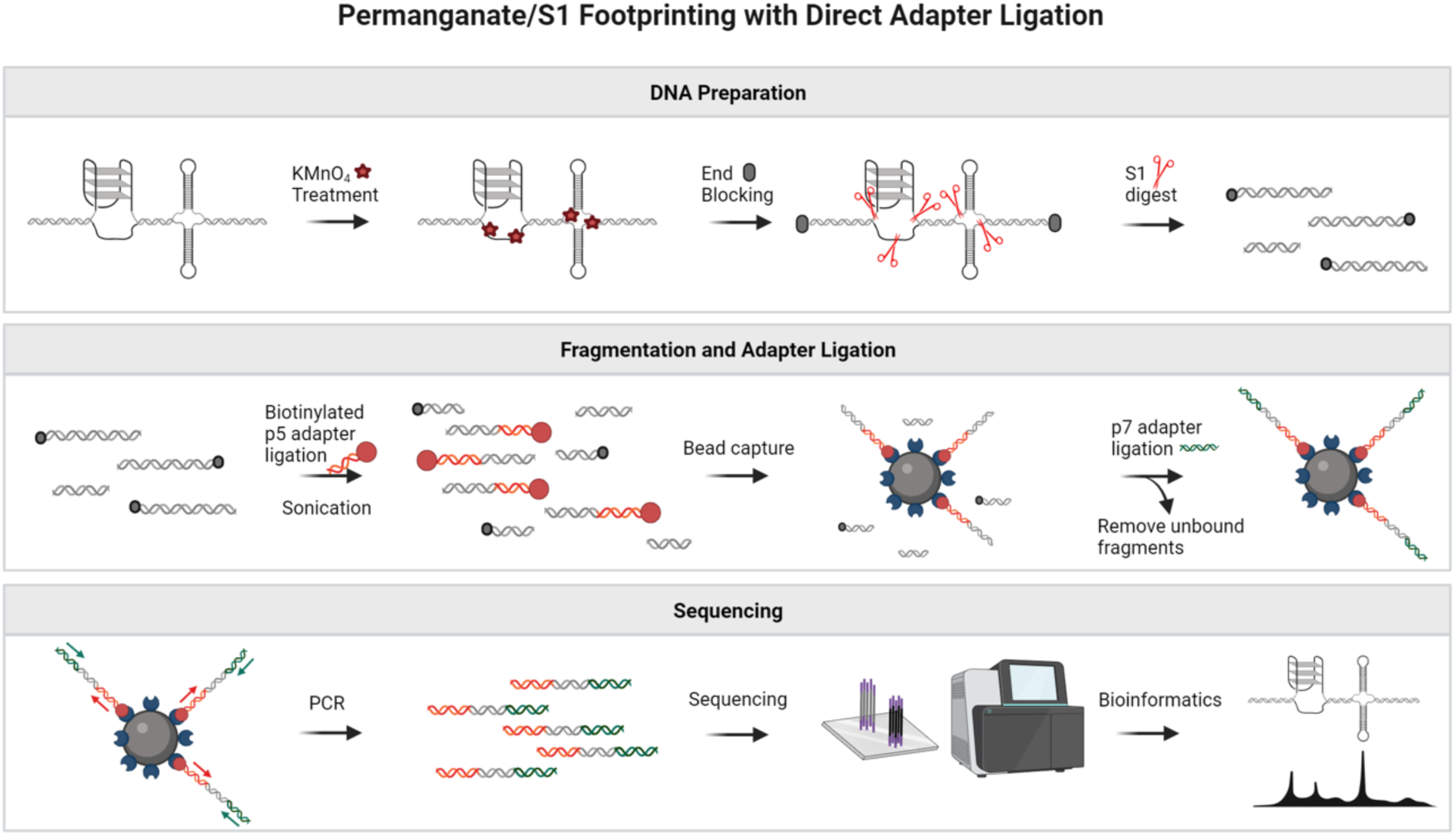
PDAL-Seq pipeline. Schematic description of each step of PDAL-Seq. Created with Biorender.com.

In PDAL-Seq, DNA is digested with S1 nuclease creating breaks at non-B DNA regions followed directly by coupling with the biotinylated Illumina P5 adapter. These biotinylated fragments are then fragmented by sonication and bound to streptavidin-coated beads. Only fragments which carry the biotinylated P5 adapter can be bound, and all others are removed during stringent washing steps. Next, the P7 adapter is ligated to the fragments attached to the beads. Finally, a PCR enriches all fragments coupled to the streptavidin beads, which results in material of sufficient quantity for Illumina sequencing. After read alignment and stringent filtering as described in the methods part, bamCoverage files are generated and visualized with the IGV tool (Robinson et al., 2011). Figure 3 shows the results of three different PDAL-Seq experiments conducted in kidney (HEK293), lung (H1299) and breast (MCF7) cell lines, respectively. In addition, we show an antibody based G4-ChIP experiment in HEK293 (C. Li et al., 2021), as well as the *in vitro* G4 formation in HEK293 cells stabilized with pyridostatin obtained via G4-Seq (Marsico et al., 2019). Here we show the promoter regions of three different genes, *PDPK1*, *SKI* and *TERT*, where G4 motifs overlap (black bars, 1. track) with *in vitro* G4-Seq (black track, 2. track) and G4-ChIP (blue, 4. track) signals compared to the input material (grey, 3. track) in HEK293 cells. We show PDAL-Seq examples for HEK293 (red, 5. track), H1299 (green, 6. track) and MCF7 cells (orange, 7. track).

**Figure 3.**
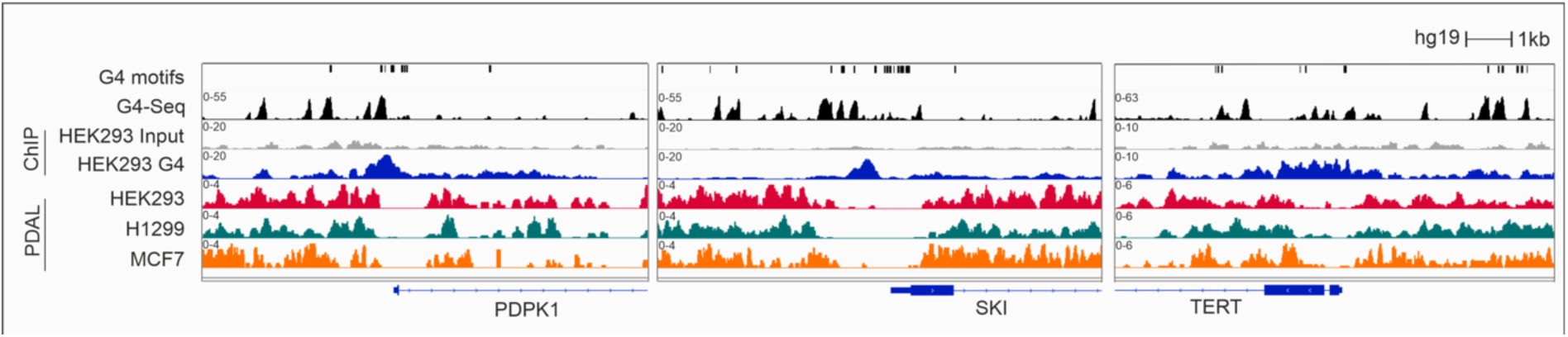
PDAL-Seq tracks showing the *PDPK1, SKI* and *TERT* loci. G4 motifs (black, top track), *in vitro* G4-Seq with PDS (black, (Marsico et al., 2019).), HEK293 input (grey) and ChIP-Seq data (blue, (C. Li et al., 2021)), as well as different PDAL-Seq experiments in HEK293 (red), H1299 (green) and MCF7 (orange) cells.

Due to the chemical approach of PDAL-Seq, G4 formation is detected as valleys, since these structures are cut out by S1 nuclease. Compared to the other methods, PDAL-Seq is much better at detecting the entire range of ssDNA formation, and not just those that align with G4 motifs.

In comparison to the original Permanganate/S1 Nuclease Footprinting method described by Kouzine et al., our protocol uses ten times less starting material and works with significantly reduced volumes of enzymes and chemicals, which results in fewer overall costs. Another major improvement is the recovery of the ssDNA regions leading to higher quality libraries and subsequent Illumina sequencing. In the original protocol, digested DNA fragments were directly biotinylated with terminal transferase followed by binding to streptavidin beads. Since the strong non-covalent binding of biotin to streptavidin requires harsh conditions to reverse, the loss of DNA fragments during this step was substantial (e.g. starting material before binding to beads 14 µg vs eluted material 70 ng). PDAL-Seq overcomes these major hurdles and allows for more efficient and in-depth characterization of non-B DNA structure formation in living cells at a more affordable cost.

## 2. Technical details: Choosing the appropriate potassium permanganate and S1 concentrations

Choosing the appropriate KMnO_4_ and S1 nuclease concentrations are critical to obtain high-quality sequencing data. Inappropriately high KMnO_4_ concentrations result in over-fragmentation of the DNA, whereas concentrations that are too low result in weak or unstable non-B DNA structures that are lost during DNA extraction. A detailed literature search revealed that a wide range of KMnO_4_ concentrations have been used for ssDNA footprinting experiments ranging from 0.5-3.5 mM (Davis, Capp, Record, & Saecker, 2005) to 40 mM (Gilchrist et al., 2012; Kouzine et al., 2017), or even 80 mM (Kouzine et al., 2019). KMnO_4_-treated DNA is highly sensitive to S1 nuclease and therefore, the KMnO_4_ concentration mainly determines the amount of S1 nuclease required for fragmentation. Several methods using S1 nuclease digestion utilized a wide range of units (U): 200 U S1 in S1-END Seq (Matos-Rodrigues et al., 2022) or 9 U S1 in S1-Seq (Mimitou & Keeney, 2018), Kouzine 0-300 U S1 (Kouzine et al., 2017). Careful optimization steps revealed that 20-40 mM KMnO_4_ in combination with 0.25-0.5 U S1 nuclease per 1 µg DNA results in an optimal fragmentation, with most of the DNA still being of high molecular weight and some degree of fragmentation ranging from 1-10 kb visible as a smear on an 0.6% agarose gel or a fragment analyzer run.

## 3. Method description

### 3.1. Material

● Pipettors and low-retention tips
● Centrifuge (refrigerated)
● Qubit fluorimeter and high sensitivity dsDNA kit (Cat # Q32851)
● Covaris M220 sonicator or equivalent device (e.g. Bioruptor)
● 130 µl Covaris sonication tubes
● Agarose gel apparatus and detection system
● PCR machine (Thermocycler)
● AMPure or SPRI beads (Beckman Coulter)
● Dynabeads kilobaseBinder kit, streptavidin-coated magnetic beads (Thermo Scientific, Cat # 60101)
● Magnetic stand
● 50 ml and 15 ml tubes
● 2 ml and 15 ml MaXtract gel tubes for phenol:chloroform extraction (Qiagen)
● 1.5 ml and 2 ml low binding tubes
● 200 µl PCR tubes

### 3.2. Reagents

● KMnO_4_ (Merck, Cat # 223468-25G)
● MilliQ ultra-pure water (do not use DEPC treated water!)
● 0.5 M EDTA
● β-Mercaptoethanol
● Proteinase K (20 mg/ml stock concentration)
● RNase A (20 mg/ml stock concentration, *New England Biolabs* Cat # T3018L)
● Phenol/Chloroform/Isoamyl alcohol (25:4:1)
● 2 M NH_4_OAc
● 100% EtOH absolute
● 1 M Tris-HCl pH 7.5 and 8.0
● 10 mM Tris-HCl pH 7.5 and 8.0
● 1× TE buffer pH 8.0 (10 mM Tris-HCl pH 8.0 and 1 mM EDTA)
● Terminal Transferase (400 U/µl, Merck Cat # 3333566001)
● 10 mM Cordycepin-5’-triphosphate (Merck Cat # C9137-1MG)
● S1 nuclease (10,000 U, Promega, Cat # M5761)
● T4 DNA ligase (*New England Biolabs,* Cat # M0202S)
● T4 DNA polymerase (*New England Biolabs,* Cat # M0203S)
● DNA End Repair Mix (*New England Biolabs* Cat # E6050)
● Phusion HSII polymerase (2 U/µl, Thermo Scientific, Cat # F549S)
● Agarose for gel electrophoresis

### 3.3. Solutions

● <u>70% EtOH:</u> Mix 7 ml of 100% EtOH abs. with 3 ml sterile filtered deionized water
● <u>250 mM KMnO</u>_4_ <u>solution:</u> Dissolve 0.79 g KMnO_4_ in 20 ml deionized water. Stir 1h at 37°C. Protect from light and prepare shortly before usage.
● <u>40 mM KMnO</u>_4_ <u>solution:</u> Mix 160 µl of 250 mM KMnO_4_ solution with 840 µl deionized water. Protect from light. Prepare shortly before usage and bring to 37°C in a water bath prior usage.
● <u>Low salt solution:</u> **Table 1.**
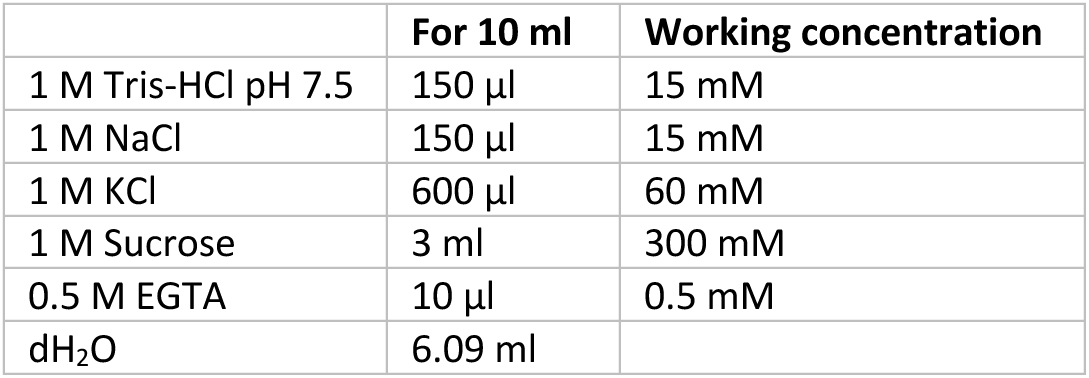
Preparation of low salt solution. Bring to 37°C in a water bath prior usage.
● <u>Stop solution:</u> **Table 2.**
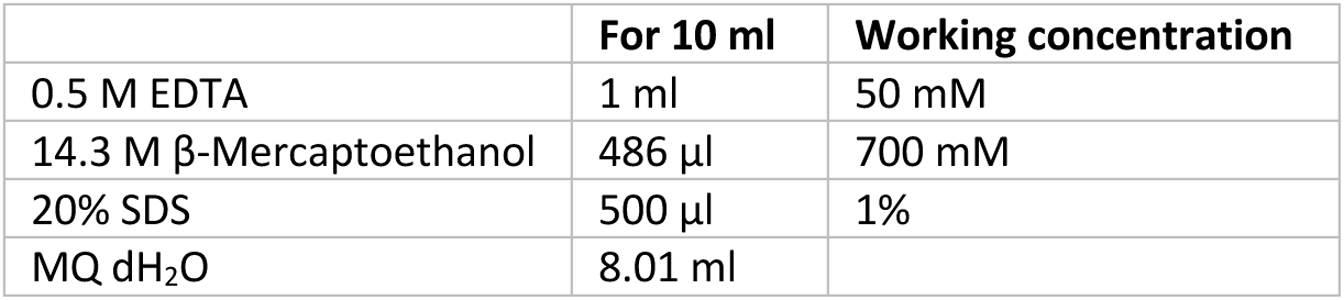
Preparation of stop solution. Protect from light. Prepare shortly before usage and bring to 37°C in a water bath prior usage.

### 3.4. Adapters and indexing primers

● P5 and P7 adapters (ordered in HPLC quality): **Table 3.**
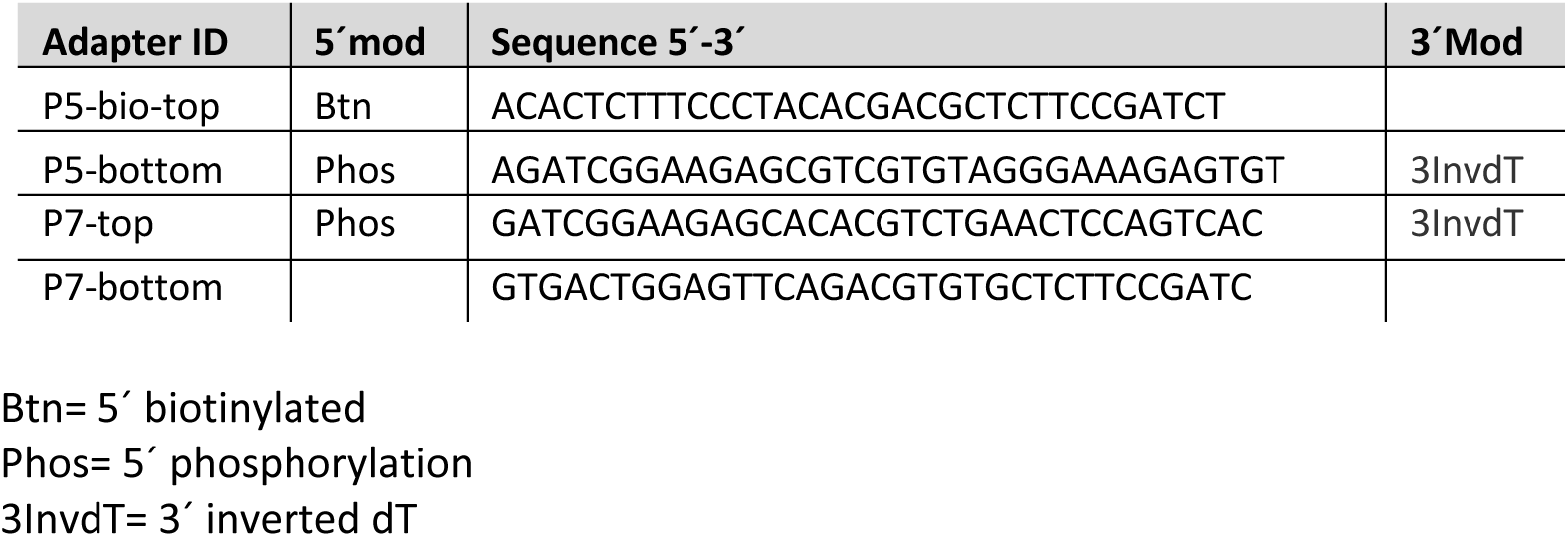
P5 (biotinylated) and P7 adapters.
● i5 and i7 dual indexing primers (*New England Biolabs* Cat # E7600S or order individual primers): **Table 4.**
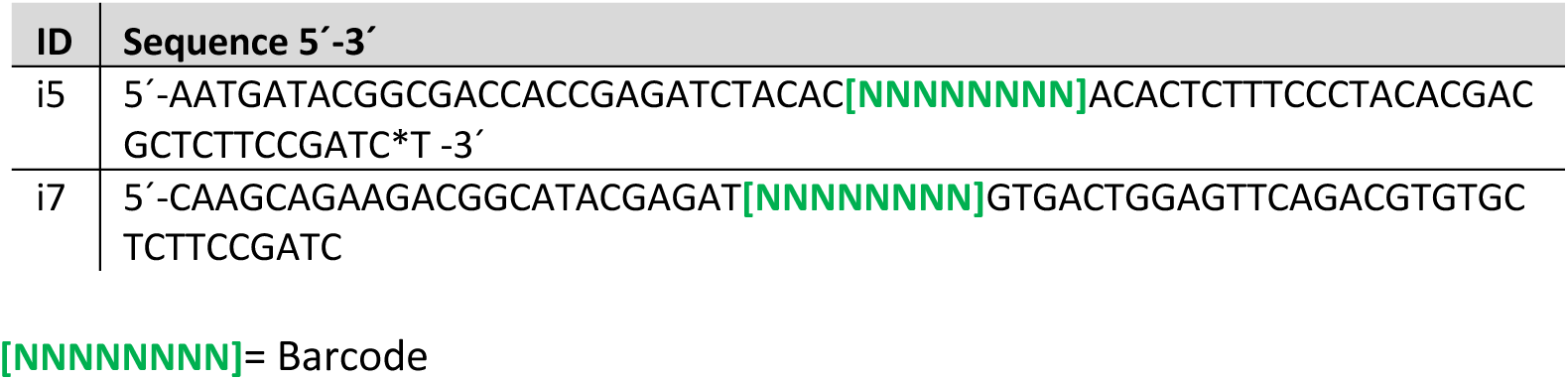
i5 and i7 dual indexing primers.

### 3.5. Timetable

The following protocol is conducted in eight days according to table 5.

**Table 5.**
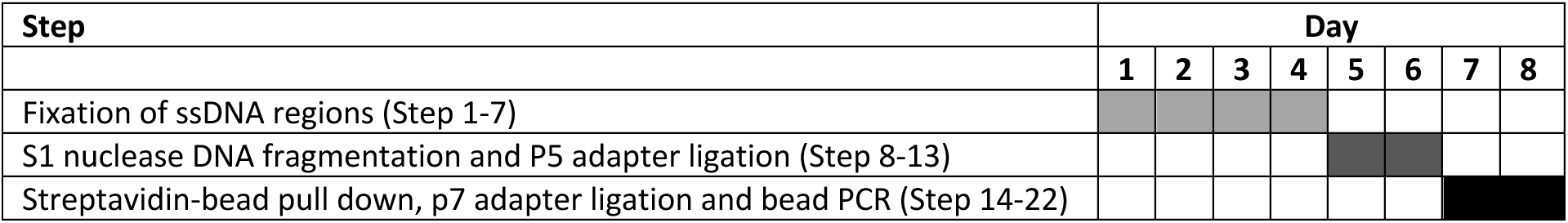
Timetable.

### 3.6. Fixation of ssDNA regions and DNA preparation (Day 1-4)

#### Step 1a: Permanganate fixation step with adherent cells (Day 1)

1. Prepare a T-75 culture flask with approx. 60-80% confluent cells containing approx.∼5-8 x 10^6^ cells.
2. Wash twice with 10 ml warm 1× PBS buffer (without Ca^2+^ and Mg^2+^).
3. Add 2 ml warm low salt buffer.
4. Gently swirl the flask and let it incubate for 1 min.
5. Add 667 µl of warm 40 mM KMnO_4_ stock solution and incubate for 80 sec. at 37°C in the incubator protected from light. (*see Note 1*)
6. Add 2.67 ml stop solution and swirl carefully until the cellular lysate gets clear.
7. Transfer the 5.33 ml lysate with a serological pipette into a 15 ml tube.
8. Add 79.5 µl of 20 mg/ml proteinase K (300 µg/ml final concentration) to digest the proteins.
9. Resuspend well by inverting at least 10 times and perform a quick spin. Avoid pipetting.
10. Incubate at 37°C overnight without shaking. (*see Note 2*)

#### Step 1b: Permanganate fixation step with suspension cells (Day 1)

1. Prepare a T-75 culture flask with approx. ∼5-8 x 10^6^ cells.
2. Centrifuge with 300xg for 5min.
3. Remove the supernatant.
4. Wash twice with 10 ml warm 1× PBS buffer (without Ca^2+^ and Mg^2+^).
5. Centrifuge with 300xg for 5min
6. Remove the supernatant.
7. Add 2 ml warm low salt buffer.
8. Gently swirl the flask and let it incubate for 1min.
9. Add 667 µl of warm 40 mM KMnO_4_ stock solution or dH_2_O and incubate for 80 sec. at 37°C in the incubator protected from light. (*see Note 1*)
10. Add 2.67 ml stop solution and swirl carefully until the cellular lysate gets clear.
11. Transfer the 5.33 ml lysate with a serological pipette into a 15 ml tube.
12. Add 79.5 µl of 20 mg/ml proteinase K (300 µg/ml final concentration) to digest the proteins.
13. Resuspend well by inverting at least 10 times and perform a quick spin. Avoid pipetting.
14. Incubate at 37°C overnight without shaking. (*see Note 2*)

#### Step 2: PCI DNA extraction (Day 2)

1. Spin a 15 ml high-density Maxtract phase-lock gel tube for 5 min at 2,000xg to pellet the gel.
2. Transfer the DNA lysate (approx. 6 ml) directly into the phase-lock gel tube.
3. Add 6 ml phenol/chloroform/isoamyl alcohol (25:24:1) solution (1 volume).
4. Invert gently at least ten times and spin down at 2,000xg for 5 min.
5. Transfer the supernatant to a new 50 ml tube.
6. Add 3 ml of 2 M NH_4_OAc (0.5 volumes) to the sample.
7. Add 18 ml ice-cold 100% ethanol (2 volumes) and mix by inverting (solution gets cloudy and you will see a clot forming). (*see Note 3*)
8. Precipitate the DNA in the –20 °C freezer for 30 min.
9. Spin at 2,000xg at 4°C for 60 min.
10. Carefully remove and discard supernatant.
11. Add 2 ml ice-cold 80% ethanol.
12. Spin with 2,000xg at 4°C for 5 min.
13. Carefully remove and discard supernatant.
14. Let tube dry inverted on a paper towel for ∼30-60 min.
15. Let tube dry upright for another ∼30-60 min. (*see Note 4*)
16. Resuspend the DNA pellet in 1 ml of 10 mM Tris-HCl buffer pH 8.0 by inverting and/or flicking the tube. Avoid pipetting.
17. Let the DNA equilibrate at room temperature until the DNA is completely dissolved. (*see Note 5*)

#### Step 3: RNase A digest (Day 3) (see Note 6)

1. Add 2.5 µl of 20 mg/ml RNase A to the DNA sample.
2. Mix by inverting and spin down. Avoid pipetting.
3. Incubate at 37°C for 1 h.

#### Step 4: PCI DNA extraction (Day 3)

1. Spin a 15 ml high-density Maxtract phase-lock gel tube for 5 min at 2,000xg to pellet the gel.
2. Transfer the DNA lysate (1 ml) directly into the phase-lock gel tube
3. Add 1 ml phenol/chloroform/isoamyl alcohol (25:24:1) solution (1 volume).
4. Invert gently at least ten times and spin down at 2,000xg for 5 min.
5. Transfer the supernatant to a new 15 ml tube.
6. Add 500 µl of 2 M NH4OAc (0.5 volumes) to 1 ml reaction volume.
7. Add 3 ml ice-cold 100% ethanol (2 volumes) and mix by inverting.
8. Precipitate the DNA in the –20°C freezer for 30 min.
9. Spin with 2,000xg at 4°C for 60 min.
10. Carefully remove and discard the supernatant.
11. Add 1 ml ice-cold 80% ethanol.
12. Spin with 2,000xg at 4°C for 5 min.
13. Carefully remove and discard the supernatant.
14. Let tube dry inverted for ∼30 min.
15. Let tube dry upright for ∼30 min.
16. Resuspend in 290 µl of 10 mM Tris-HCl pH 8.0 buffer and let it equilibrate at room temperature until the DNA is completely dissolved.
17. Transfer the solution to a 1.5 ml low binding tube.

#### Step 5: Free 3’ end blocking (Day 4) (see Note 7)

1. Set up the following reaction (Table 6): **Table 6.**
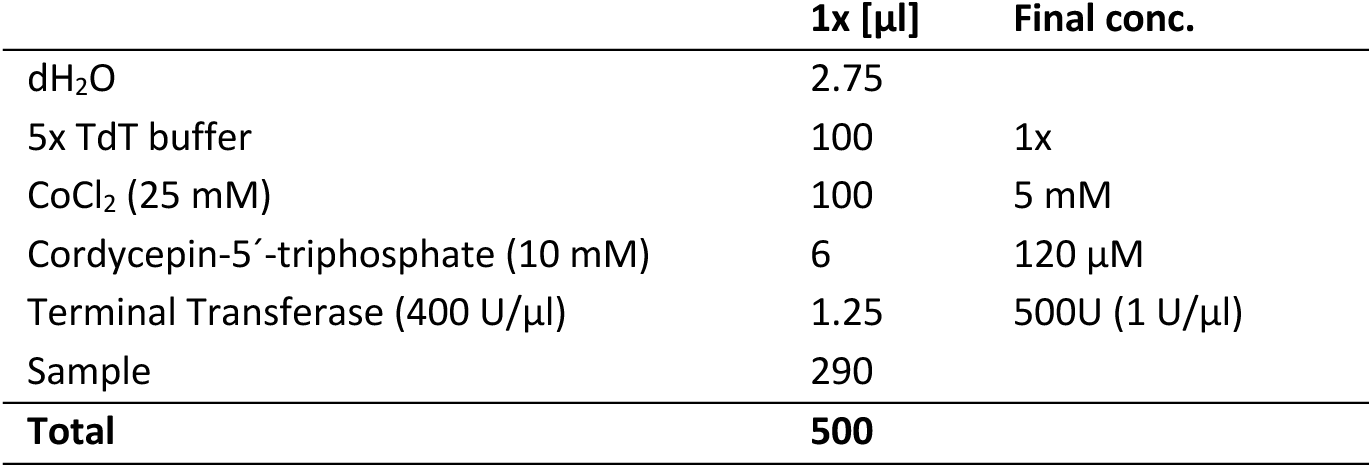
Free 3’end blocking mastermix using terminal transferase.
2. Incubate at 37°C for 1 h.

#### Step 6: PCI DNA extraction (Day 4)

1. Spin a 15 ml high-density Maxtract phase-lock gel tube for 5 min at 2,000xg to pellet the gel.
2. Transfer the DNA lysate (500 µl) directly into the phase-lock gel tube.
3. Add 500 µl phenol/chloroform/isoamyl alcohol (25:24:1) solution (1 volume).
4. Invert gently at least ten times and spin down at 2,000xg for 5 min.
5. Transfer the supernatant to a new 50ml tube.
6. Add 250 µl of 2 M NH4OAc (0.5 volumes) to 500 µl reaction volume.
7. Add 1 µl Glycogen solution. (*see Note 8*)
8. Add 1500 µl ice-cold 100% ethanol (2 volumes) and mix by inverting.
9. Precipitate the DNA at –20°C for 30 min.
10. Spin with 2,000xg at 4°C for 60 min.
11. Carefully remove and discard supernatant.
12. Add 0.5 ml ice-cold 80% ethanol.
13. Spin with 2,000xg at 4°C for 5 min.
14. Carefully remove and discard supernatant.
15. Let tube dry inverted for ∼15 min.
16. Let tube dry upright for ∼15 min.
17. Resuspend the DNA pellet in 500 µl of 10 mM Tris-HCl buffer pH 8.0 by inverting and flicking the tube. Avoid pipetting.

#### Step 7: Second DNA precipitation (Day 4) (see Note 9)

1. Add 250 µl of 2M NH4OAc (0.5 volumes) to 500 µl reaction volume.
2. Add 1 µl Glycogen solution.
3. Add 1500 µl ice-cold 100% ethanol (2 volumes) and mix by inverting.
4. Precipitate the DNA at –20°C for 30 min.
5. Spin with 2,000xg at 4°C for 60 min.
6. Carefully remove and discard supernatant.
7. Add 0.5 ml ice-cold 80% ethanol.
8. Spin with 2,000xg at 4°C for 5 min.
9. Carefully remove and discard supernatant.
10. Let tube dry inverted for ∼15 min.
11. Let tube dry upright for ∼15 min.
12. Resuspend the DNA pellet in 500 µl of 10 mM Tris-HCl buffer pH 8.0 by inverting. Avoid pipetting.
13. Let it equilibrate for 1 h at room temperature. (*see Note 10*) 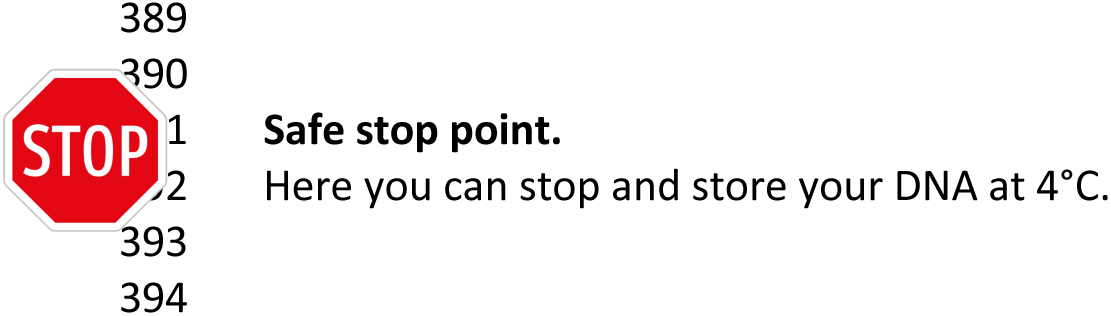

### 3.7. S1 nuclease DNA fragmentation and P5 adapter ligation (Day 5-6)

#### Step 8: S1 nuclease digest (Day 5) (see Note 11 and Note 12)

1. Measure the DNA concentration in triplicates of 1:50 dilutions with a Qubit fluorimeter and calculate the mean of each triplicate.
2. Prepare 500 µl sample dilution containing 40 µg DNA. Work on ice.
3. Prepare the mastermix given in Table 7 to digest the sample with 0.5 U S1 nuclease per 1 µg DNA. (*see Note 13*) **Table 7.**
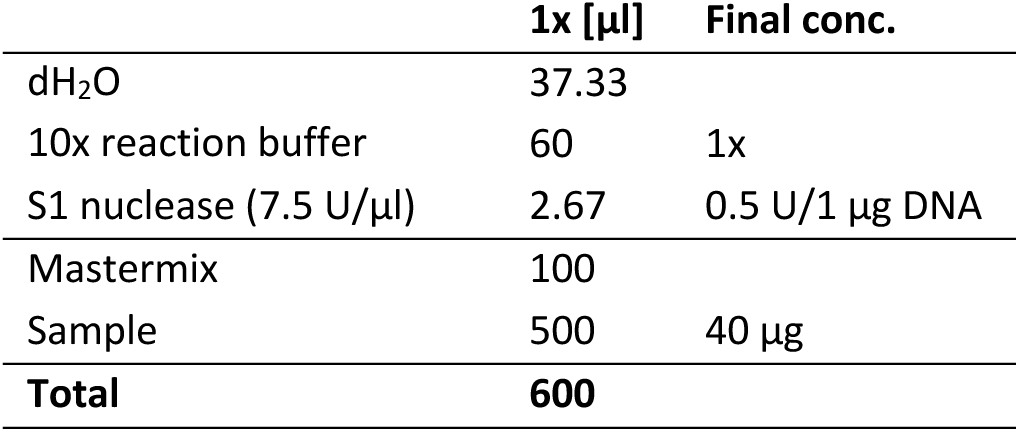
S1 nuclease digest mastermix.
4. Incubate the reactions at 37°C for 20 min.
5. Add 12 µl of 0.5 M EDTA to a final conc. of 10 mM EDTA.
6. Check your sample on a 0.6% agarose gel. Most of the DNA should be undigested, but a faint smear with a range of 2-10 kb is visible. (*see Note 14*).
7. Immediately proceed with DNA extraction.

#### Step 9: PCI DNA extraction (Day 5)

1. Spin a 15 ml high-density Maxtract phase-lock gel tube for 5 min at 2,000xg to pellet the gel.
2. Transfer the digested DNA (600 µl) directly into the phase-lock gel tube.
3. Add 600 µl phenol/chloroform/isoamyl alcohol (25:24:1) solution (1 volume).
4. Invert gently at least ten times and spin down at 2,000xg for 5 min.
5. Transfer the supernatant to a new 15 ml tube.
6. Add 300 µl of 2 M NH4OAc (0.5 volumes) to 600 µl reaction volume.
7. Add 1 µl Glycogen solution.
8. Add 1800 µl ice-cold 100% ethanol (2 volumes) and mix by inverting (solution gets cloudy).
9. Precipitate the DNA at –20°C for 30 min.
10. Spin with 2,000xg at 4°C for 60 min.
11. Carefully remove and discard supernatant.
12. Add 500 µl ice-cold 80% ethanol.
13. Spin with 2,000xg at 4°C for 5 min.
14. Carefully remove and discard supernatant.
15. Let tube dry inverted for ∼15 min.
16. Let tube dry upright for ∼15 min.
17. Resuspend in 84 µl of 10 mM Tris-HCl buffer pH 8.0.
18. Prepare a 200 µl PCR tube with 4 µl 10 mM Tris HCl buffer pH 8.0 and add 1 µl sample (1:5 dilution). Mark it as C1 for a Fragmentanalyzer/Bioanalyzer measurement.

#### Step 10: DNA end repair with T4 DNA polymerase (Day 5)

1. Check the concentration of a 1:50 dilution (in triplicates) of the samples with the Qubit HS dsDNA assay.
2. Calculate the amount of T4 DNA Polymerase, which you need. For 1 µg DNA, 1 U T4 DNA polymerase is required. (*see Note 15*)
3. Mix the following reaction on ice (Table 8): **Table 8.**
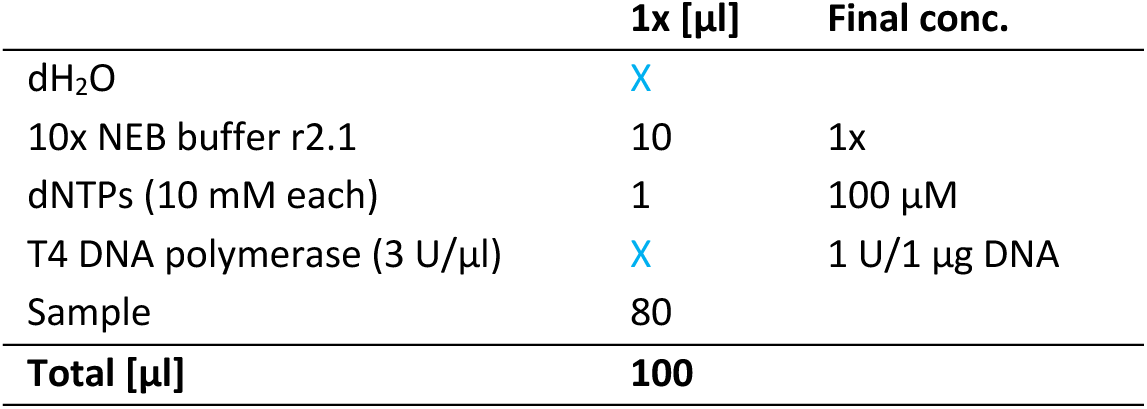
DNA end repair mastermix using T4 DNA polymerase.
4. Incubate for 15 min at 12°C in a thermocycler.
5. Add 2 µl EDTA (final conc. 10 mM) and heat inactivate at 75°C for 20 min.

#### Step 11: PCI DNA extraction (Day 5)

1. Add 100 µl Tris HCl buffer pH 8.0 to the DNA sample of Step 10.5. (*see Note 16*)
2. Spin a 1.5ml high-density Maxtract phase-lock gel tube for 5 min at 12,000xg to pellet the gel.
3. Transfer the DNA (200 µl) directly to the phase-lock gel tube.
4. Add 200 µl phenol/chloroform/isoamyl alcohol (25:24:1) solution (1 volume).
5. Invert gently at least times and spin down at 12,000xg for 5 min.
6. Transfer the aqueous phase on the gel to a new 2 ml tube.
7. Add 100 µl of 2 M NH_4_OAc (0.5 volumes) to 200 µl reaction volume.
8. Add 1 µl Glycogen solution.
9. Add 600 µl ice-cold 100% ethanol (2 volumes) and mix by inverting (solution gets cloudy).
10. Precipitate the DNA at –20°C for 30 min.
11. Spin with 12,000xg at 4°C for 30 min.
12. Carefully remove and discard supernatant.
13. Add 500 µl ice-cold 80% ethanol.
14. Spin with 12,000xg at 4°C for 5 min.
15. Carefully remove and discard supernatant.
16. Let tube dry inverted for ∼15 min.
17. Let tube dry upright for ∼15 min.
18. Resuspend in 80 µl of 10 mM Tris-HCl pH 8.0 buffer. 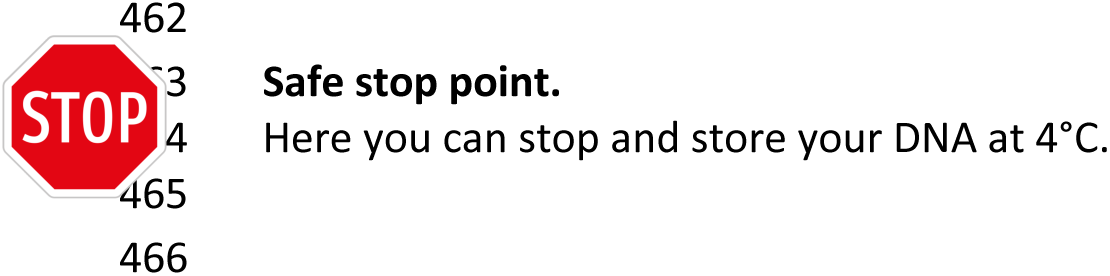

#### Step 12: P5 Adapter preparation (Day 6)

1. Mix the P5-top (biotinylated, 100 μM, Table 3) and P5-bottom (100 μM, Table 3) oligos at equimolar concentration to reach a 50 µM each concentration in 0.2 ml PCR tubes (Table 9). **Table 9.**
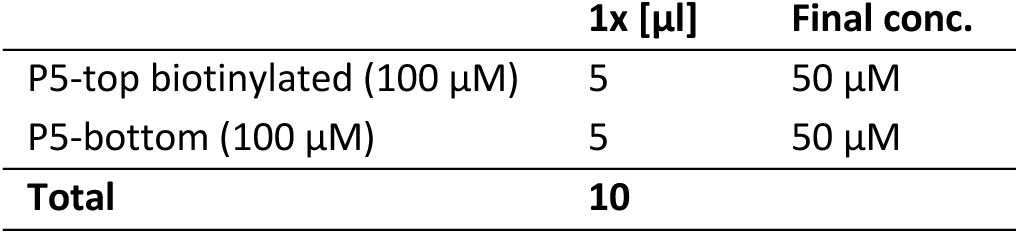
Preparation of double stranded biotinylated P5 adapters.
2. Place the tubes in a PCR machine.
3. Run the following incubation program (Table 10): (*see Note 17*) **Table 10.**
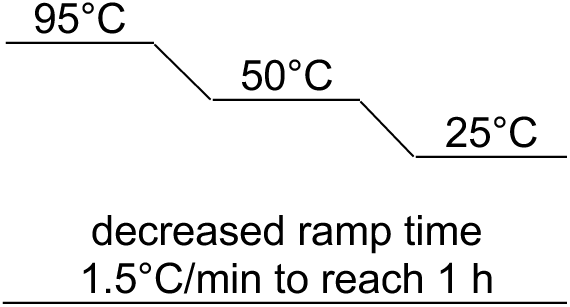
Incubation conditions for adapter preparation.

#### Step 13: P5 adapter ligation (Day 6)

1. Prepare the following T4 DNA ligase mix (Table 11) (*see Note 18*): **Table 11.**
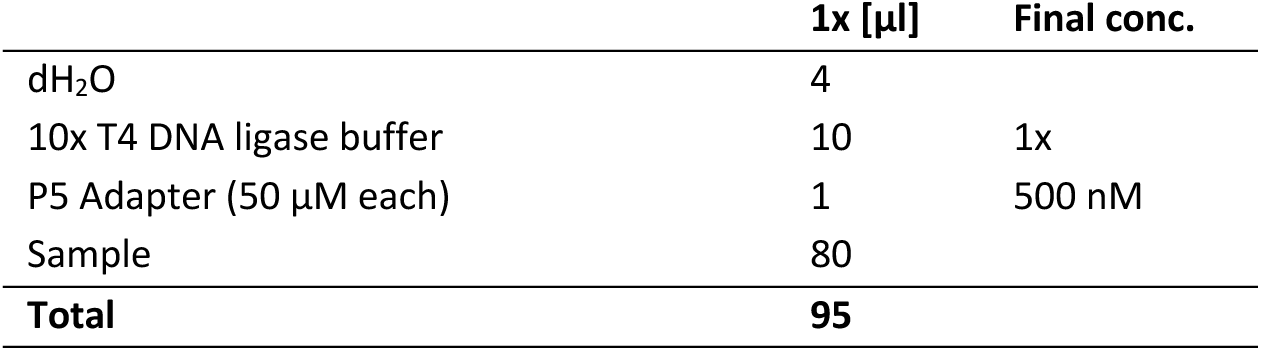
T4 DNA ligase mastermix for ligation of the biotinylated P5 adapter.
2. Incubate for 5 min on ice. (*see Note 19*)
3. Add 5 µl of 400 U/µl T4 DNA ligase (final conc 2000 U).
4. Incubate at 16°C overnight (>18 h) in a thermocycler.
5. Prepare a 200 µl PCR tube with 4 µl 10 mM Tris-HCl buffer pH 8.0 and add 1 µl sample (1:5 dilution). Mark it as C2 for a Fragmentanalyzer/Bioanalyzer measurement.

### 3.8. Streptavidin-bead pull down, P7 adapter ligation and bead PCR (Day 7-8)

#### Step 14: Purification with AMPure XP or SPRI beads (0.9 volumes ∼ ≥ 100 bp) (Day 7) **(see Note 20)**

1. Bring the beads to room temperature at least 30 min before you start.
2. Vortex the Agencourt AMPure XP/SPRI bead bottle to completely resuspend any magnetic particles that may have settled.
3. Add 100 µl of 10 mM Tris-HCl buffer pH 8.0 to your sample to get a total sample volume of 200 µl. (*see Note 21*)
4. Add 180 µl of well-mixed beads to 200 µl sample (0.9 volumes beads). Gently pipette up and down 10 times to mix thoroughly.
5. Perform a quick spin.
6. Incubate for 15 min at room temperature.
7. Prepare fresh 80% ethanol. (*see Note 22*)
8. Place the tubes on the magnetic stand to separate beads from the solution. Visually confirm that the beads have moved to the side of the tube, and the solution is clear.
9. Discard the supernatant.
10. With the tubes still on the magnetic stand, dispense 400 μl of freshly prepared 80% ethanol to each tube.
11. Remove the tube from the magnetic stand and resuspend by flicking.
12. Put tubes on the magnet, allow beads to separate, aspirate out ethanol and discard.
13. Repeat the washing steps (10-12).
14. Use a 10 µl pipette to remove all traces of ethanol.
15. Let the beads dry up to 5 min at room temp on the magnetic stand. (*see Note 23*)
16. Keeping the tube on the magnetic stand, add 132 μl of 1× TE buffer pH 8.0 to each tube, remove the tubes from the magnetic stand, and completely resuspend beads.
17. Incubate the tube at room temp for 5 min.
18. Put the tubes on the magnet for 5 min (or until solution gets clear) and transfer 132 µl supernatant to new tubes. Discard the beads.
19. Prepare a 200 µl PCR tube with 4 µl of 10 mM Tris-HCl buffer pH 8.0 and add 1 µl sample (1:5 dilution). Mark it as C3 for a Bioanalyzer measurement.

#### Step 15: DNA sonication (Day7)

1. Transfer 130 µl sample into a Covaris microTube-AFA Fiber Pre-Slit Snap Cap (Cat # 520045)
2. Sonicate the DNA for to yield an average fragment size of 400 bp.
  a. Start the PC of the Covaris.
  b. Turn on Covaris.
  c. Start the SonoLab Software.
  d. Fill in water into the water bath until water level turns green.
  e. Set the following parameters: 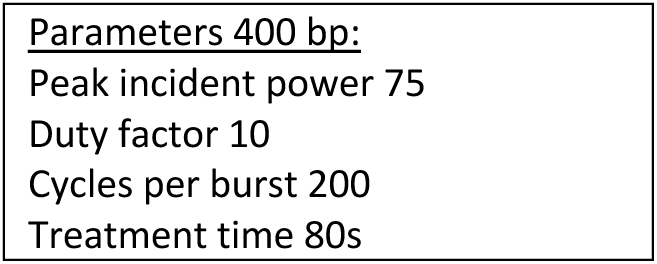
  f. Fill your sample into a microfiber tube.
  g. Place the tube in the tube holder.
  h. Fix the tube with the upper metal arm.
  i. Close the door.
  j. Start the program.
  k. When the program is finished, remove your tube.
  l. Remove the water with the Covaris syringe. Discard the water.
  m. Wipe out the water bath with the cotton swab carefully! Do not leave back any liquid!
  n. Close the door.
3. As a quick control step, you can check the size on a 1% agarose gel. The size range should be between 100-1000 bp with a mean peak at 400 bp. (*see Note 24*)
4. Prepare a 200 µl PCR tube with 4 µl of 10 mM Tris-HCl buffer pH 8.0 and add 1 µl sample (1:5 dilution). Mark it as C4 for a Bioanalyzer measurement.

#### Step 16: Pull down step with streptavidin-coated beads (Day 7)

1. Thoroughly resuspend the Dynabeads® M-280 (kilobaseBINDER kit) in the vial (vortex >30 sec or tilt and rotate for 5 min).
2. Transfer 50 μl resuspended beads to a 1.5 ml low binding tube.
3. Place the tube on the magnet for 2 min.
4. Carefully remove and discard the supernatant while the tube remains on the magnet. Avoid touching the bead pellet with the pipette tip.
5. Add 200 μl binding solution (included in the kit) along the inside wall of the tube where the beads are collected.
6. Gently resuspend by pipetting.
7. Place the tube on the magnet for 2 min and remove the supernatant.
8. Resuspend the beads in 200 μl binding solution.
9. Add the solution containing the biotinylated DNA-fragments to the resuspended beads. Mix carefully.
10. Incubate the tube at room temperature for 1.5 h on a wheel tube rotor with about 10 rpm to keep the beads in suspension.
11. Place the tube on the magnet and remove the supernatant.
12. Wash the Dynabeads®/DNA-complex twice in 500 μl washing solution (included in the kit).
13. Wash 2× with 400 µl of 10mM Tris-HCl buffer pH 7.5.
14. Keep the beads in 400 µl of 10 mM Tris-HCl pH 7.5 at 4°C until you proceed.

#### Step 17: End Repair (Day 7)

1. Mix the following reaction on ice (Table 12): **Table 12.**
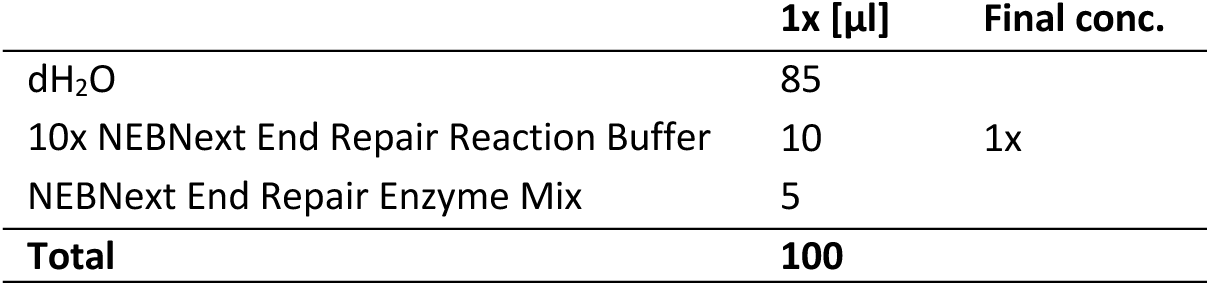
NEBNext End Repair Mastermix preparation.
2. Place the tube on the magnet and remove the supernatant.
3. Wash once with 400 µl 10mM Tris-HCl buffer pH 7.5.
4. Place the tube on the magnet and remove the supernatant.
5. Resuspend the beads in 100 µl end repair mix and transfer the bead mix to a 200 µl PCR tube.
6. Incubate in a thermocycler for 30 min at 20°C with heated lid set to 30°C.
7. Wash the beads three times with 200 µl of 10 mM Tris-HCl buffer pH 7.5.
8. Keep the beads in 200 µl of 10mM Tris-HCl pH 7.5 until you proceed.

#### Step 18: P7 adapter preparation (Day 8)

1. Mix the P7-top (100 μM, Table 3) and P7-bottom (100 μM, Table 3) oligos at equimolar concentration to reach a 50 µM each concentration in 0.2 ml PCR tubes (Table 13). **Table 13.**
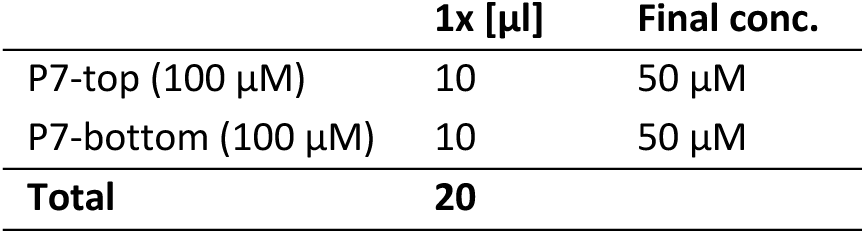
Preparation of double stranded P7 adapters.
2. Place the tubes in a PCR machine.
3. Run the following incubation program (Table 14). (*see Note 17*) **Table 14.**
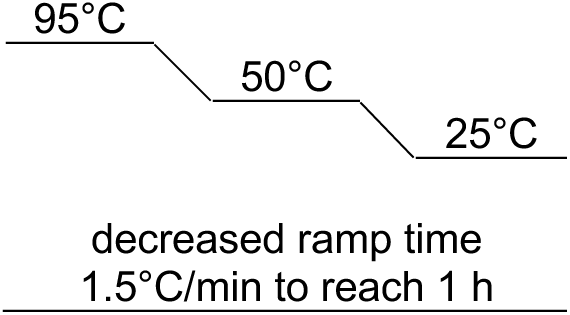
Incubation conditions for adapter preparation.

#### Step 19: P7 adapter ligation (Day 8)

1. Prepare the following T4 DNA ligase mix on ice (Table 15): **Table 15.**
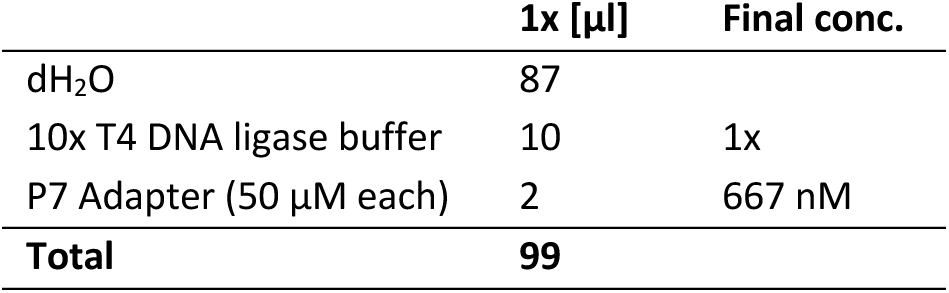
T4 DNA ligase mastermix for ligation of the P7 adapter.
9. Resuspend the beads in 99 µl T4 ligase mix.
10. Incubate for 5 min on ice.
11. Add 1 µl 400 U/µl T4 ligase (final conc. 400 U) and mix well by inverting.
12. Incubate at 20°C for 4 h on a wheel tube rotator.
13. Wash the beads 2× with 200 µl washing buffer.
14. Wash the beads 2× with 200 µl of 10 mM Tris-HCl buffer pH 7.5. 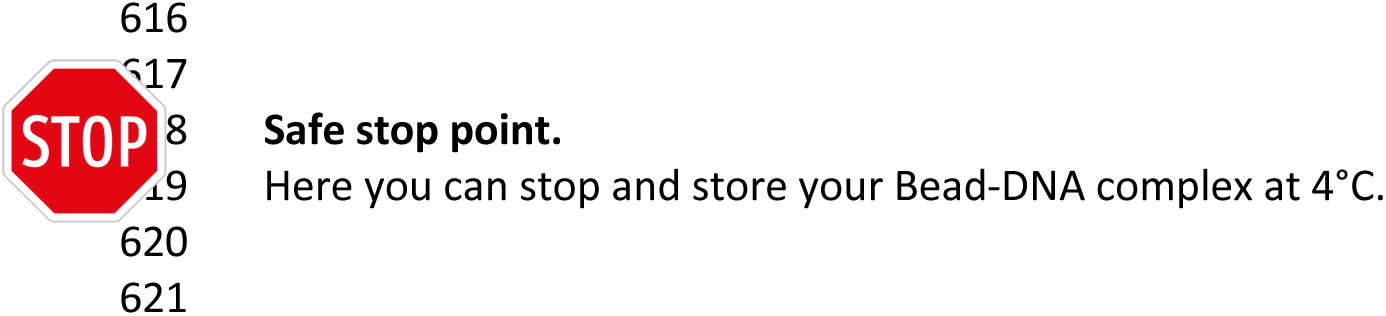

#### Step 20: PCR (Day 8)

The combination of PCR primers (indexes, Table 4) is important and depend also on the number of samples which you want to combine. Guidelines how dual indexing primers are combined can be found directly at the web page of Illumina (Illumina, 2023).

1. Prepare the following PCR reaction (Table 16): **Table 16.**
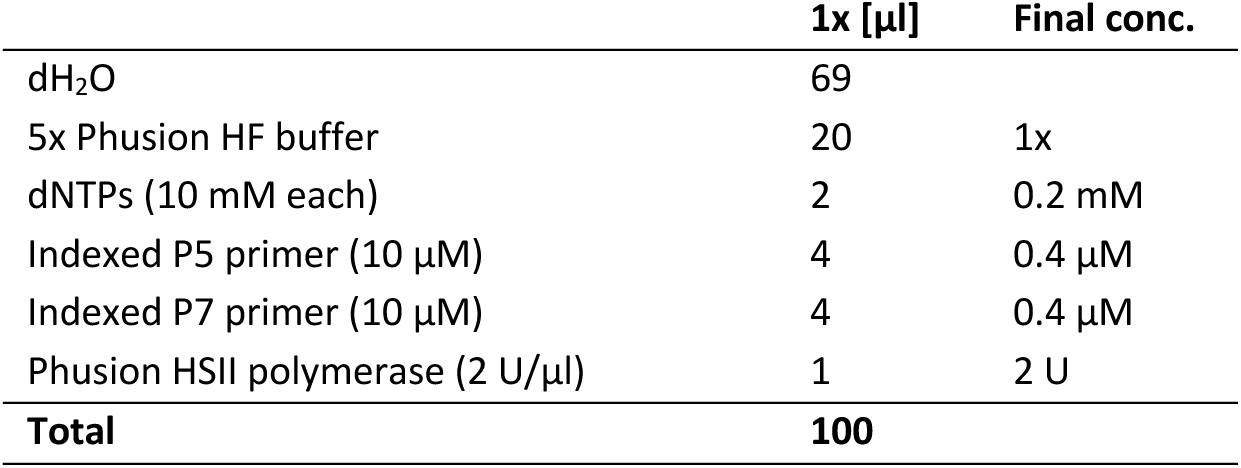
Indexing PCR mastermix using Phusion HSII polymerase.
2. Place the beads on a magnetic stand and remove the Tris-HCl pH 7.5 buffer.
3. Resuspend the beads in 100 µl PCR mix.
4. Run the PCR with the following conditions (Table 17): **Table 17.**
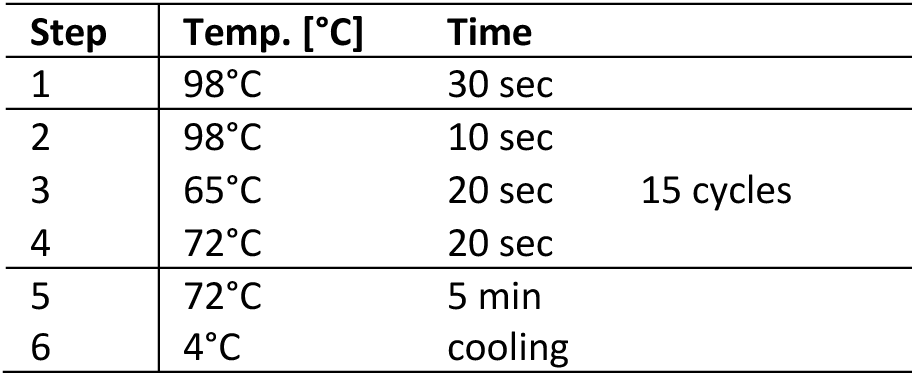
PCR cycling conditions for indexing PCR. After 15 min (∼6 cycle) interrupt the PCR and pipette up and down 2-3× to resuspend the beads again. (*see Note 25*)
5. Capture the beads on the magnetic stand.
6. Transfer the supernatant (∼90 µl) into a fresh 1.5 ml low binding tube.

#### Step 21: Purification with AMPure XP OR SPRI beads (0.9 volumes ∼ ≥ 200 bp) (Day 8)

1. Gently shake the Agencourt AMPure XP/SPRI bead bottle to completely resuspend any magnetic particles that may have settled. The beads must be at room temperature before use.
2. Add 81 µl of well-mixed beads to 90 µl sample (0.9 volumes beads). Gently pipette up and down 10 times to mix thoroughly.
3. Perform a quick spin.
4. Incubate for 15 min at room temperature.
5. Prepare fresh 80% ethanol.
6. Place the tubes on the magnetic stand to separate beads from the solution. Visually confirm that the beads have moved to the side of the tube, and the solution is clear.
7. Discard the supernatant.
8. With the tubes still on the magnetic stand, dispense 200 μl of freshly prepared 80% ethanol to each tube.
9. Remove the tube from the magnetic stand and resuspend by flicking.
10. Put tubes on the magnet, allow beads to separate, aspirate out ethanol and discard.
11. Repeat the washing steps (8-10).
12. Use a 10 µl pipette to remove all traces of ethanol.
13. Let the beads dry up to 5 min at room temp on the magnetic stand.
14. Keeping the tube on the magnetic stand, add 21 μl of 10 mM Tris-HCl buffer pH 8.0 to each tube, remove the tubes from the magnetic stand, and completely resuspend beads.
15. Incubate the tube at room temp for 5 min.
16. Put the tubes on the magnet for 5 min (or until solution gets clear) and transfer 20 µl supernatant to new tubes. Discard the beads.
17. Prepare a 200 µl PCR tube and add 1 µl sample. Mark it as C5 for a Fragmentanalyzer/ Bioanalyzer measurement.
18. Quantify 1 µl (undiluted) with Qubit HS dsDNA assay.

#### Step 22: Quality control, DNA concentration measurement and sequencing (Day8)

1. Load the samples marked with C1-C5 on a Bioanalyzer Chip/Fragmentanalyzer.
2. If the C5 sample does not show any adapter contamination, they are ready for sequencing.
3. Determine the concentration of the library either with Qubit Fluorimetric dsDNA method or with a qPCR-based detection method.
4. Sequencing is performed on NextSeq or NovaSeq with at least 50 M reads per sample. Initial experiments were performed with single end sequencing, latest runs were performed with paired end sequencing.

## Bioinformatic analysis

The obtained raw reads are checked for quality and aligned to the human reference genome using BWA-MEM (H. Li & Durbin, 2009). Non-unique reads with a mapping quality less than 30 are excluded with SAM tools (Danecek et al., 2021). After stringent filtering excluding reads overlapping ENCODE blacklisted regions or low mappability regions, coverage files (bamCoverage, bin size 10-50) are visualized in IGV tools (Robinson et al., 2011). Optionally, non-B formation can be strand-specific visualized by choosing the option forward or reverse in “*Only include reads originating from fragments from the forward or reverse strand*” during the generation of bamCoverage files. Default is *none* to not separate by strands. This pipeline is available at https://usegalaxy.org/u/kaivan/h/pdal-seq-collection-1. In order to map the non-B DNA regions revealed by PDAL-seq in a high throughput manner, a “valley finder” is employed and the results are overlapped with computationally predicted non-B sequence motifs. This pipeline is available at https://github.com/kxk302/MACS2. Test data generated from HEK293 and Raji cells are available at https://usegalaxy.org/u/kaivan/h/pdal-seq-data.

There are many existing tools that automatically detect peaks in ChIP-seq data, but they do not detect valleys (i.e. areas of the genome with no coverage flanked by areas with high coverage). Therefore, in order to use one of the existing peak callers, the valleys in coverage files must be converted into peaks. To that end, pyBigWig is used (accessible by https://github.com/deeptools/pyBigWig) to convert bigWig coverage files into a tabular format appropriate for histograms (chromosome, interval, number of reads). Then, for each chromosome in a histogram file, the number of reads is deducted from the maximum number of reads. This flips the data in such a way that valleys become peaks. Then, Model-based Analysis of ChIP-Seq (MACS2) (Zhang et al., 2008) is used as a general peak caller. MACS2 is run for various window sizes (25, 35, 50, 75, 100, 150, 200) to find peaks. MACS2’s output is then converted into the bed format that contains peak intervals for each chromosome.

For detecting non-B DNA regions, gfa (accessible by https://github.com/abcsFrederick/non-B_gfa; a suite of programs developed at NCI-Frederick/Frederick National Lab to find sequences associated with non-B DNA forming motifs) is used to detect Short Tandem Repeats (STR), Direct Repeats (DR), Mirror Repeats (MR), and Inverted Repeats (IR). Quadron (Sahakyan et al., 2017) is used to detect G4 motifs. Quadron (and not gfa) is used for G4 detection as Quadron is specifically tailored for G4 detection. A-Phased Repeats (APR) or Z DNA motifs are disregarded since PDAL-seq would not detect them, as these structures do not form single-stranded DNA. gfa and Quadron’s output are converted into bed format containing non-B DNA intervals for each chromosome.

The intersect utility of bedtools (Quinlan & Hall, 2010) is used to find the intersection between peaks and non-B DNA annotations for each chromosome. In order to determine the best parameters for valley detection, a quality control step is performed. Briefly, a summary of the intersects is created by dividing the sum of the length of intersect intervals by the sum of the length of peak (sequencing valleys or valleys) intervals for each chromosome. For each chromosome, this ratio objectively measures how well the sequencing valleys intersect with non-B DNA annotations. In our experience, valleys with window width set to 25 bp best overlap with non-B DNA annotations. The overlap ratio of all chromosomes is averaged using harmonic mean to find the overall overlap ratio which can be translated as the percentage of non-B DNA motifs which are overlapping with valleys in the sequencing data. We found with our data the best overlap ratio was 81%. Comparing these ratios with a ratio created using random intervals from the genome (not containing non-B forming motifs), it was found that non-B motifs overall have roughly a 10-fold higher overlap with valleys than randomly generated intervals throughout the genome overlap with valleys. Furthermore, the non-B motifs also show the same elevated intersect ratio when overlapping with valleys than the ratio of non-B motifs overlapping with randomly generated intervals throughout the genome (Fig. 4).

**Figure 4.**
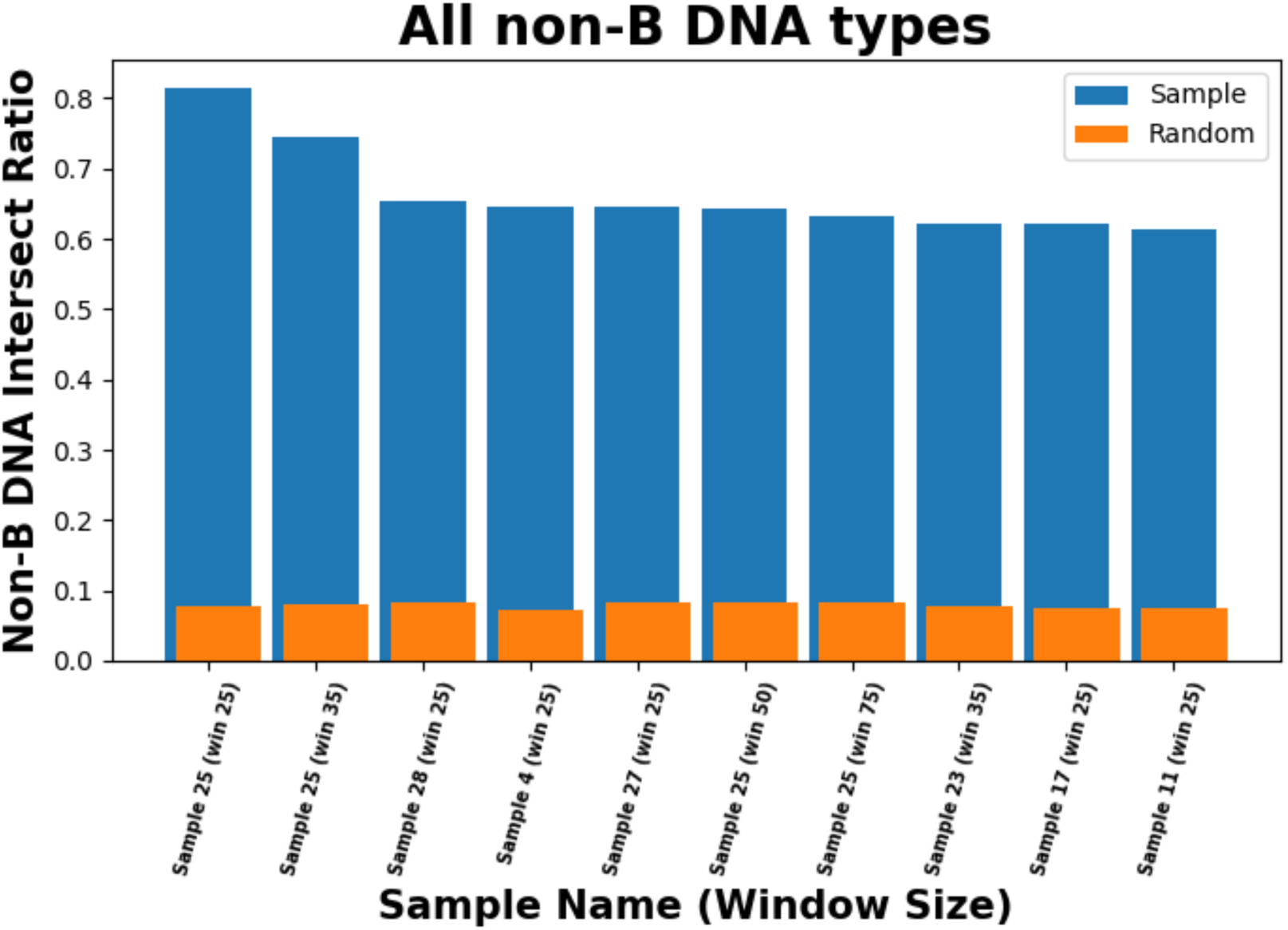
Non-B forming motif intersection ratios with sequencing valley intervals. For each sample, we calculate the intersect ratio between PDAL-seq’s valley intervals (blue bars) and non-B DNA annotation intervals and compare it to intersect ratio between random intervals and non-B DNA annotation intervals (orange intervals). We observe that PDAL-seq’s valley intervals intersect with non-B DNA annotations nearly an order of magnitude more than random intervals

## Notes

● ***Note 1*:** Prepare low salt buffer, stop solution and KMnO_4_ solutions approximately 1 h before you start the experiment and bring it to 37°C in a water bath. Furthermore, a control reaction with 0 mM KMnO_4_ can be included. Especially for the first trials we would recommend that, since the evaluation of the fragmentation degree is easier with a control sample. The 0 mM KMnO_4_ sample is highly insensitive for S1 nuclease, except in very high concentrations (Fig. 5).
● ***Note 2*:** The overnight incubation step can be performed in a 37°C water bath, if you do not have a thermo block for 15 ml tubes.
● ***Note 3*:** Mix until the phase separation has disappeared. Otherwise, you observe a transparent clot after the DNA extraction, which cannot be resolved anymore.
● ***Note 4:*** The DNA pellet changes the color during drying from white to clear.
● ***Note 5*:** Depending on the DNA concentration complete equilibration could take over night.
● ***Note 6:*** The RNA digest step is done after the first DNA extraction, since you would need very high RNase A concentrations in the initial lysis buffer of Step 1.
● ***Note 7*:** This step is important to block all DSBs and SSBs present in the DNA so that later only breaks resulting from non-B DNA structures can be detected.
● ***Note 8:*** We use glycogen solution to get a better visible DNA pellet.
● ***Note 9:*** The second DNA precipitation step should help to get rid of any residual Cordyceptin in the sample, which could interfere with later steps.
● ***Note 10*:** Wait until the DNA is completely dissolved or increase the volume, otherwise concentration measurements will not be correct and the S1 digest will be insufficient.
● ***Note 11:*** This step requires optimization. Try this reaction with only 1 µg DNA in 15 µl reaction volume and try different concentrations of S1 nuclease. We experienced that S1 nucleases of different companies have different digestion capacities. Most of the DNA should not be digested and only a small smear until about 100 bp should be visible (Fig. 5). Over-digestion leads to many false positive signals after sequencing.
● ***Note 12*:** For the final run, we start with 40 µg DNA in the library preparation, although there will be still material left. Keep it as backup in the fridge if something goes wrong and repeat the steps. Especially for the first trials.
● ***Note 13*:** The S1 nuclease (as many restriction enzymes) have different activities at different DNA concentrations. So, keep volumes constant. We optimized it for 1 µg in 15 µl digestion volume and scaled it then up to 40 µg in 600 µg (factor 0.067 µg/µl)
● ***Note 14*:** We suggest checking the DNA on a 0.6% agarose gel. Most of the DNA should not be digested (Fig.5). Just a faint smear is allowed. Otherwise, you see a high fragmentation degree during sequencing data analysis.
● ***Note 15:*** This step is **<u>CRITICAL!</u>** Too much T4 DNA polymerase shortens the DNA ends, due to the exonuclease activity of the enzyme!
● ***Note 16:*** We add some additional 10 mM Tris-HCl buffer pH 8.0 to elevate the extraction volume. You can choose the volume, which is comfortable for you. We experienced that the extraction works better in higher volumes.
● ***Note 17:*** Alternatively, you can heat up the PCR cycler to 95°C for 5 min with a heated lid, then turn off the cycler and let it cool down slowly for at least 1 h to allow adapter renaturation.
● ***Note 18:*** Thaw the T4 ligase buffer and the annealed biotinylated P5 adapters on ice.
● ***Note 19:*** This preincubation step without T4 DNA ligase on ice improves ligation efficiency.
● ***Note 20:*** It is critical to remove all free P5 adapters. Otherwise, they will compete with the long biotin-labeled DNA fragments in the binding with the streptavidin beads and saturate them so that you lose your DNA. You even can perform a bioanalyzer/fragmentanalyzer/tape station run prior the streptavidin pull down step to make sure all free adapters have been removed (Fig. 6).
● ***Note 21:*** We increase the sample volume here to get an efficient cleanup. In case that you use a 0 mM sample, low volumes will result in insufficient binding of the DNA sample to the beads. We experienced a great loss of DNA, if we start with too low volumes.
● ***Note 22:*** In general, we prepare fresh 80% ethanol for bead cleanups, since too low ethanol concentrations lead to elution of DNA from the beads during the washing steps.
● ***Note 23:*** Do not over dry the beads, otherwise the DNA elution will be inefficient and results in decreased recovery. Elute the samples when the beads are still dark brown and glossy looking, but when all visible liquid has evaporated. When the beads turn lighter brown and start to crack they are too dry.
● ***Note 24:*** Figure 7 shows an example for sonication range of 500 ng DNA loaded on a 0.6% agarose gel and size distribution on the Agilent Fragmentanalyzer.
● ***Note 25*:** From now on work in a post PCR workspace to avoid cross contaminations of adapters and PCR primers.

**Figure 5.**
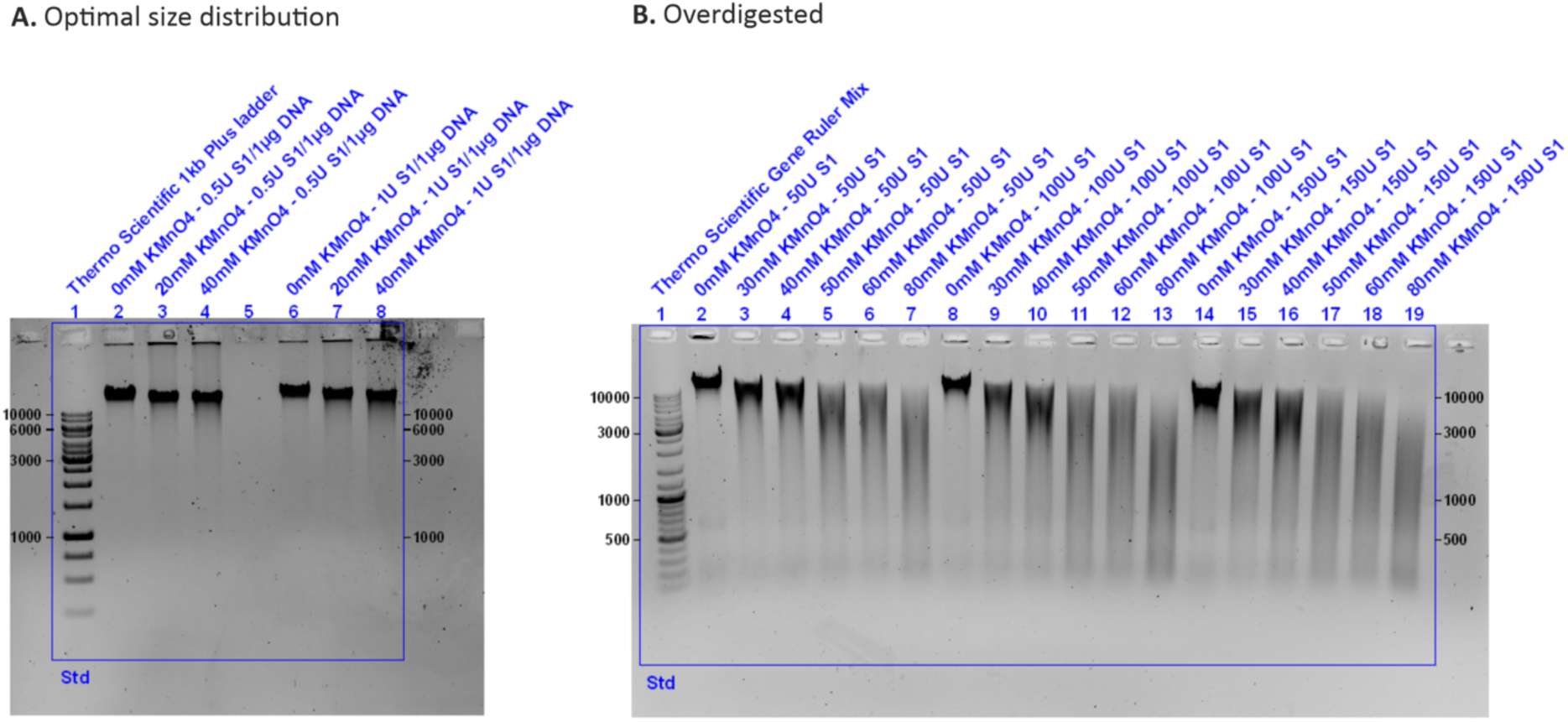
Size distribution with different KMnO_4_ and S1 nuclease concentrations. **A.** shows an optimal fragment size, where most of the DNA is still undigested, but a faint smear is visible until ∼2 kb. **B**. Shows completely overdigested DNA, which results in a high false positive rate after sequencing. The 0 mM KMnO**_4_** control is insensitive for S1 nuclease degradation, except for very high S1 nuclease concentrations such as 150 U (gel B, lane 14).

**Figure 6.**
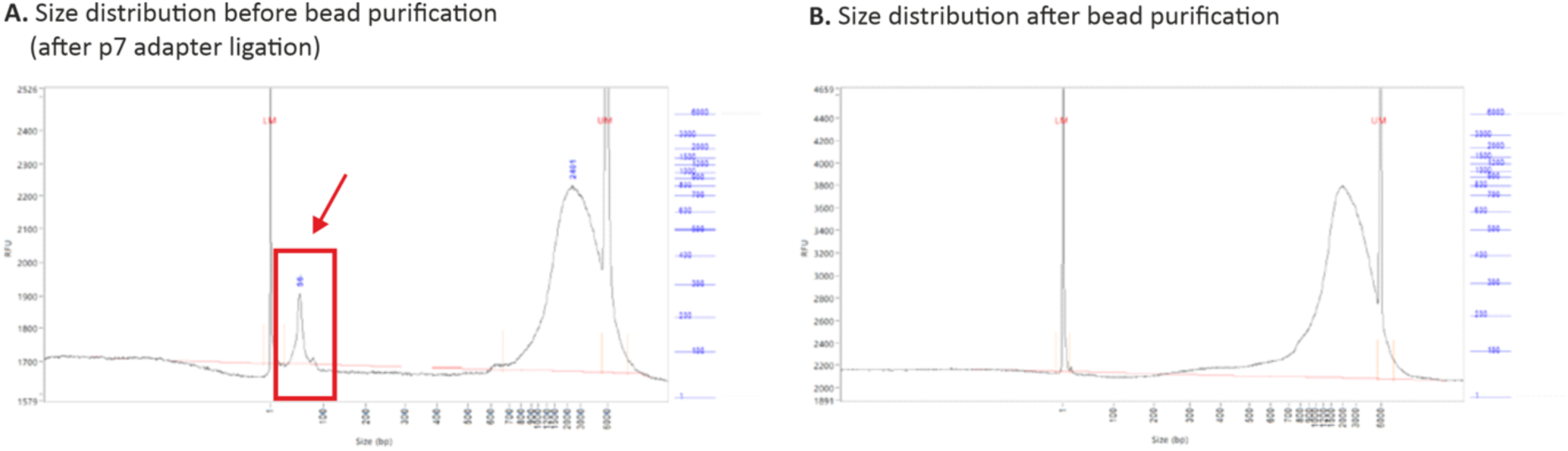
Size distribution during library preparation steps. The fragment size was determined on an Agilent Fragmentanalyzer 12 capillary system. **A.** shows the size distribution directly after the P7 adapter ligation with unligated P7 adapters left (red box and red arrow). **B.** An optimal size distribution is shown after SPRI bead clean up without any unligated P7 adapters.

**Figure 7.**
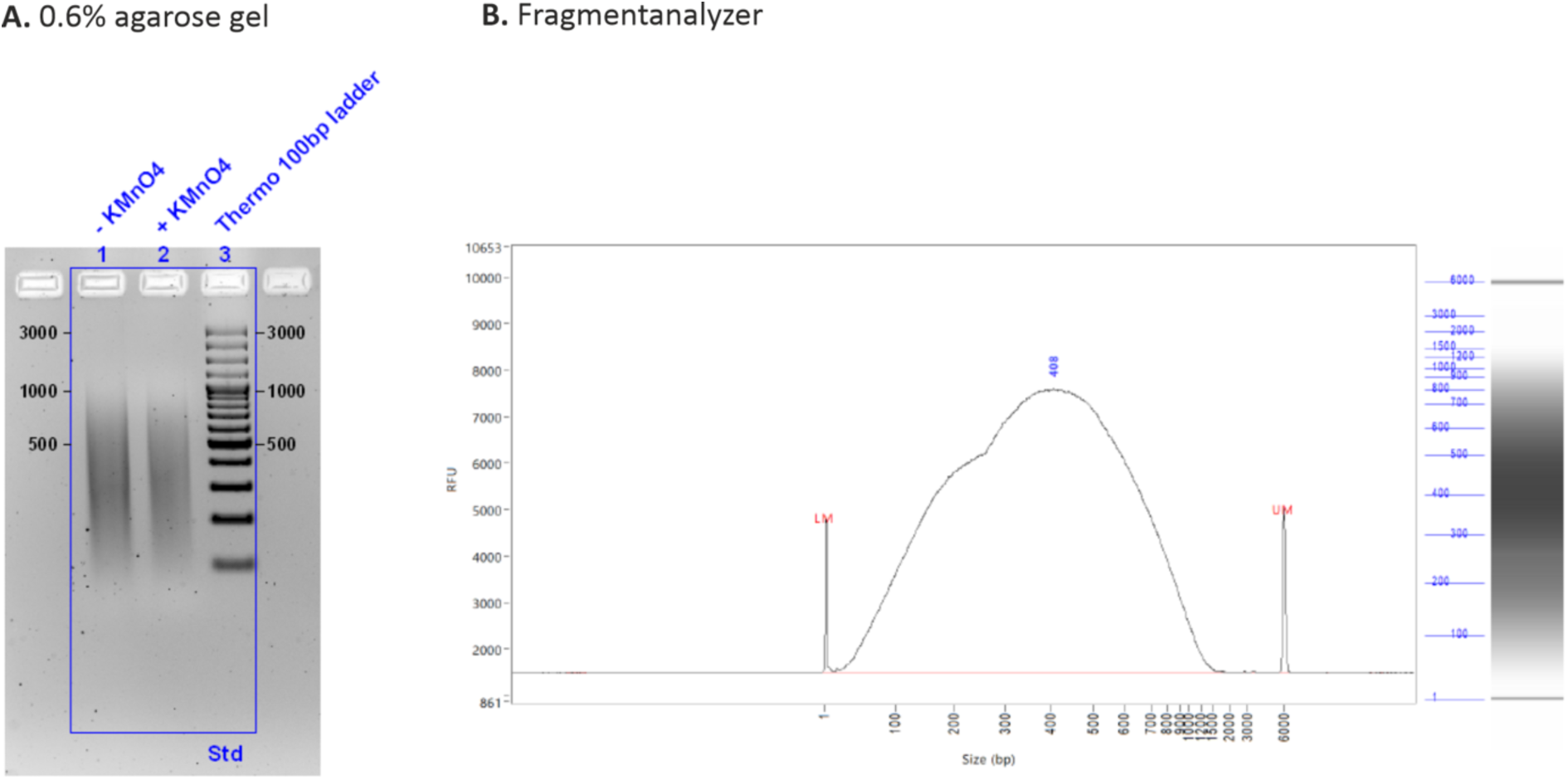
Fragment size distribution. **A.** A representative 0.6% agarose gel with 500 ng DNA loaded is shown. **B.** Control sample marked with C4 analyzed on an Agilent Fragmentanalyzer with a HS DNA kit.

